# Direct N-Me Aziridination Reaction Enables Pinpointing C=C Bonds in Lipids with Mass Spectrometry

**DOI:** 10.1101/2022.04.24.489320

**Authors:** Guifang Feng, Ming Gao, Rongrong Fu, Qiongqiong Wan, Tianze Wang, Zhourui Zhang, Suming Chen

## Abstract

The biological functions of lipids largely depend on their chemical structures. The position of C=C bonds is an essential attribute that determines the structures of unsaturated lipids. Here, we developed a new type of chemical derivatization method for C=C bond using aziridination reaction. This new cyclization method for the C=C bonds in lipids based on the direct N-Me aziridination reaction of olefins using *N*-methyl-*O*-tosylhydroxylamine (TsONHCH_3_) as the aminating reagent. When combined with the tandem MS analysis, this novel activation approach for C=C bonds enables the accurate identification their positions in different kinds of unsaturated lipids. Furthermore, an integrated workflow has been established for comprehensively identifying the C=C bond positional isomers of lipids in complicated biological sample. This work provided a new chemical approach for the structural lipidomics.

## Introduction

Lipids are an important class of bioactive molecules that are not only important components of cell membranes, but also play important roles in energy storage and cellular signal transduction of living organisms.^[1]^ The structural heterogeneity of different classes of lipids empowers their diverse biological functions,^[2]^ especially the abundant lipid isomers,^[3]^ ascribed from the different locations and configurations of C=Cs in fatty acyl chains.^[4]^ For example, C=C positional isomer ratio of unsaturated lipids shows significant correlations to the onset/progression of breast cancer,^[4b,5]^ and the brain ischemia/reperfusion. Lipid acyl chain *cis*-C=C bond position was also found to modulate plasma membrane domain registration/anti-registration.^[6]^ Therefore, the characterization of detailed structures of lipid isomers is the prerequisite to understand their biological roles.

Mass spectrometry (MS) has become the preferred method for lipid isomer analysis, due to its advantages in high sensitivity and structural resolving power.^[7]^ To address the increasingly recognized challenge in structure elucidation of complex C=C bond isomeric lipids,^[8]^ the chemically derivative methods were developed to provide access to alternative fragmentation channels and structural information.^[9]^ The cyclization of C=C bond is a desirable strategy that could not only provide rapid derivation to complex unsaturated lipids but also generate unambiguous fragments denoting the C=Cs locations in tandem MS analysis.^[3]^ So far, almost all the widely used cyclization reactions for lipid C=C position identification form the oxygen-containing heterocycles products, such as ozonides,^[9a,10]^ epoxides,^[11]^ and oxetanes.^[9c,9d,12]^ Development of a novel activation method of C=C bond in lipid would extend the toolbox of its location identification.

Aziridination of alkenes is a classic reaction for the synthesis of aziridines. The reaction was proceeded by transferring nitrenes to C=C bond typically by means of transition metal catalysts.^[13]^ Recently, a highly efficient Rh(II)-catalyzed aziridination reaction was developed for the direct preparation of unactivated aziridines from olefins using *O*-(sulfonyl)hydroxylamines as the aminating agent.^[14]^ The reactions proceed with a high stereospecificity. We reason that the aziridination could also be extended to unsaturated lipids, and form aziridine structure at the C=C positions. The ring-strain associated with aziridine may provide access to position-specific fragmentation in MS/MS to denote the location of C=Cs in fatty acyl chains of lipids. In addition, the formed *N*-methylaziridine structure possesses high proton affinity (934.8 kJ/mol)^[15]^ that would greatly enhance the sensitivity for the identification of unsaturated lipids.

In this study, we developed a new type of cyclization method for the C=C bonds in lipids based on the direct N-Me aziridination reaction of olefins using *N*-methyl-*O*-tosylhydroxylamine (TsONHCH_3_) as the aminating reagent. When combined with the tandem MS analysis, this novel activation approach for C=C bonds enables the accurate identification their positions in different kinds of unsaturated lipids. Furthermore, an integrated workflow has been established for comprehensively identifying the C=C bond positional isomers of lipids in complicated biological sample. This work provided a new chemical approach for the structural lipidomics.

## Results and discussion

### Validation of the aziridination reaction with unsaturated fatty acids

To verify our hypotheses, we first tried to develop an optimal reaction system for the aziridination of unsaturated lipids. The reactions were optimized using oleic acid [FA 18:1 (9*Z*)] as the model substrate in the presence of *N*-methyl-*O*-tosylhydroxylamine (TsONHCH_3_) as the aminating reagent, and chelating dirhodium (II) Rh_2_(esp)_2_ as the catalyst. A reaction time of only 5 min was needed to achieve an aziridination yield of more than 90% with 10 mol% Rh_2_(esp)_2_ as the catalyst in 2,2,2-trifluoroethanol (TFE) solution (Figure 2a).

**Figure 1.**
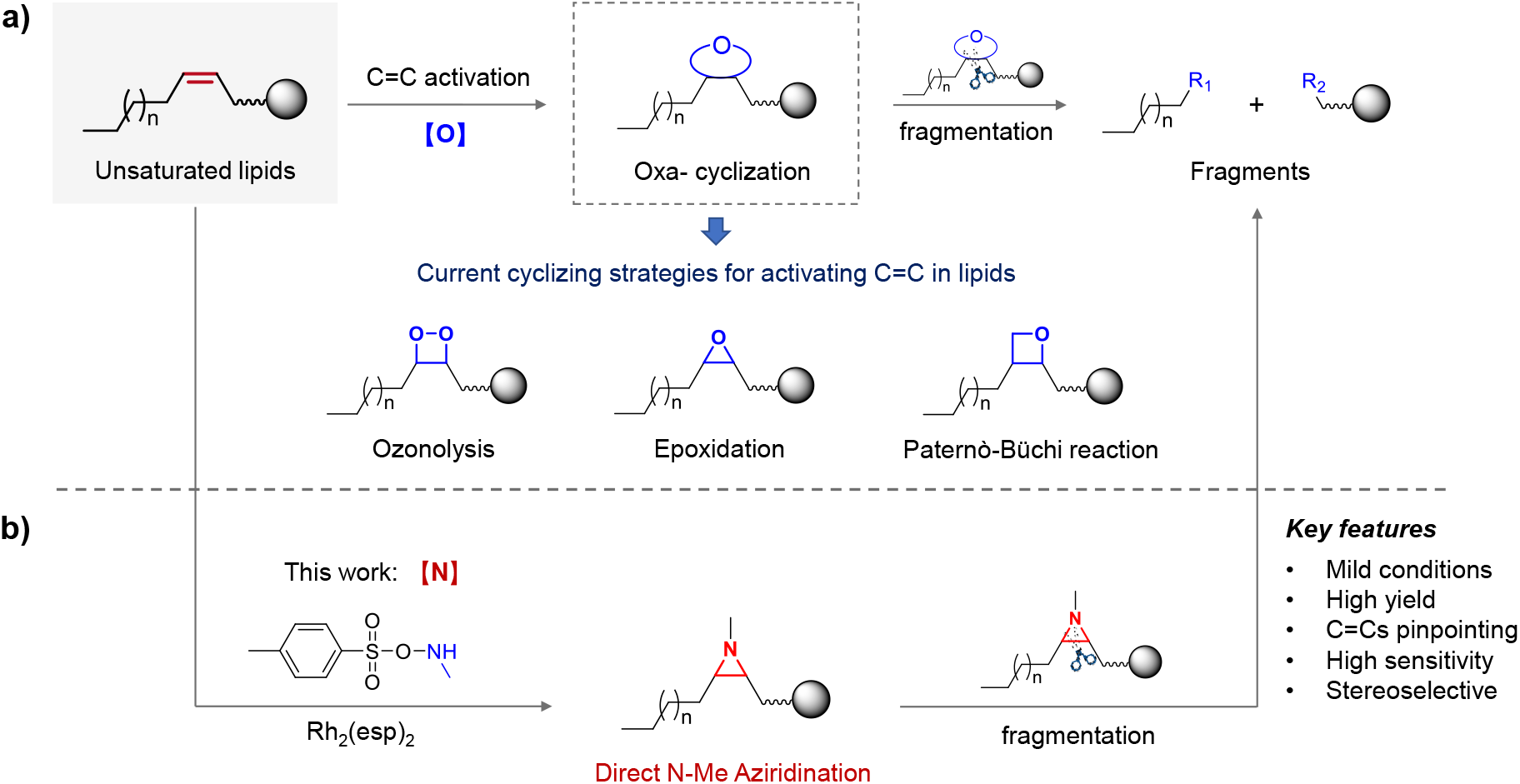
Strategies of cyclization activation of C=C bond in unsaturated lipids. **a)** The current representative cyclizing strategies for activating C=C bond in lipids that form oxygen-containing heterocycles. **b)** The proposed direct N-Me aziridination strategy for pointing C=C bond in this study.

**Figure 2.**
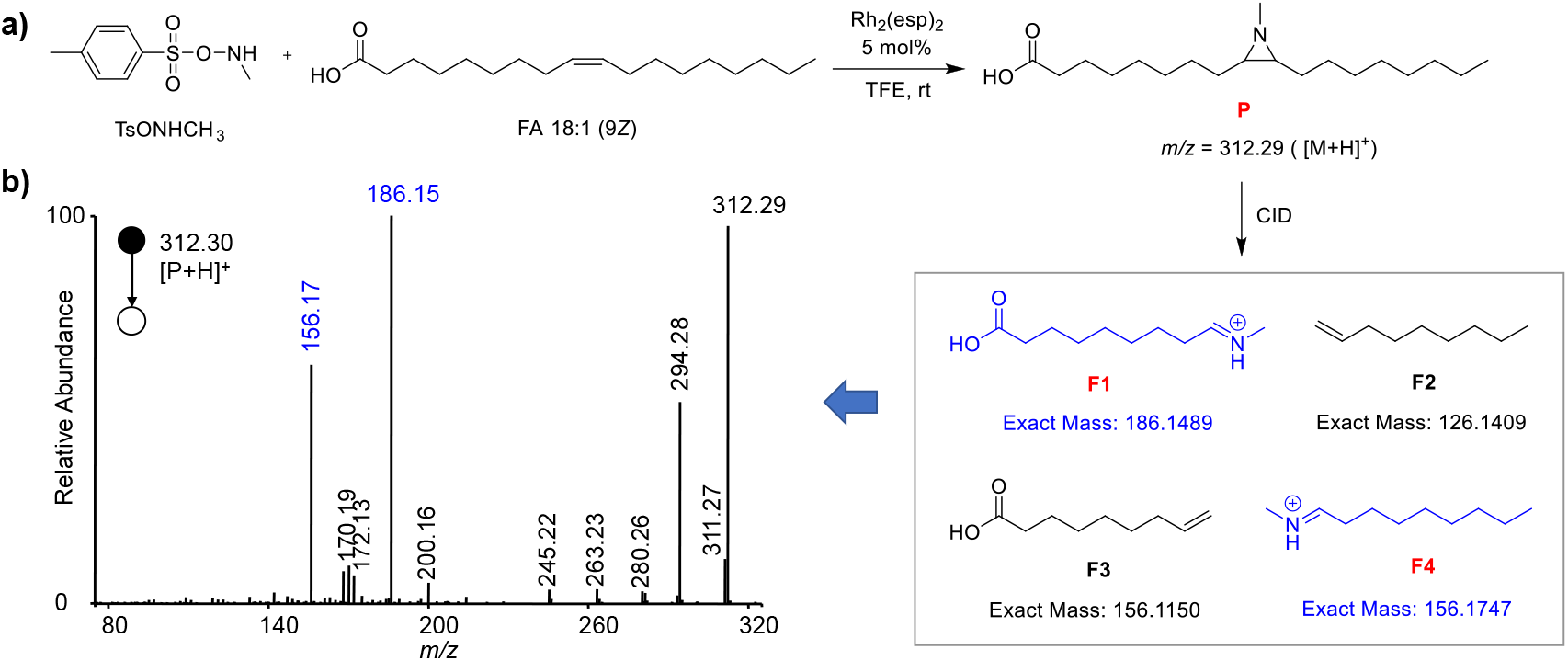
The aziridination reaction for locating the C=C bond in FA 18:1. **a)** The reaction of TsONHCH_3_ with FA18:1 (9*Z*) and the fragmentation pathway of the product ion in collision induced dissociation. b) MS/MS spectrum of the production ion (**P**) at *m/z* 312.30.

Having established the photocatalytic aziridination reaction system, we next examined the MS/MS behavior of the formed aziridine derivative to test the capability for identifying the C=C positional isomers. We speculate that there would be two types of fragmentation in this aziridine structure (Figure 2a). One way is to break from the carbon-carbon bond and the left (carboxy-terminated) carbon-nitrogen bond, resulting in fragments containing carboxyl and nitrogen positive ions (**F1**), and neutral loss of the alkyl carbon chain (**F2**). The other way is to break from the carbon-carbon bond and the right (carbon-terminated) carbon-nitrogen bond, resulting in neutral loss of carboxyl and alkyl-containing chains (**F3**), as well as fragment of alkyl chains containing nitrogen positive ions (**F4**). The MS/MS spectrum of the protonated aziridination product of oleic acid at *m/z* 321.29 ([M+H]^+^) generated by collision induced dissociation (CID) confirmed this speculation (Figure 2b). Two of the nitrogen positive ion fragments **F1** and **F4** were detected evidently at *m/z* 186.15 and 156.17, respectively, which could be used as a pair of characteristic ions for denoting the position of C=C bond. Similarly, other monounsaturated fatty acids were found to have high reactivities with TsONHCH_3_ under the same reaction conditions. The MS/MS spectra of the aziridination products of FA 18:1 (6*Z*), FA 18:1 (11*Z*), FA 18:1 (9*Z*) and FA 18:1 (9*E*) were shown in Figure 3. All the corresponding characteristic ions indicating the positions of C=C bond in fatty acids were detected clearly.

**Figure 3.**
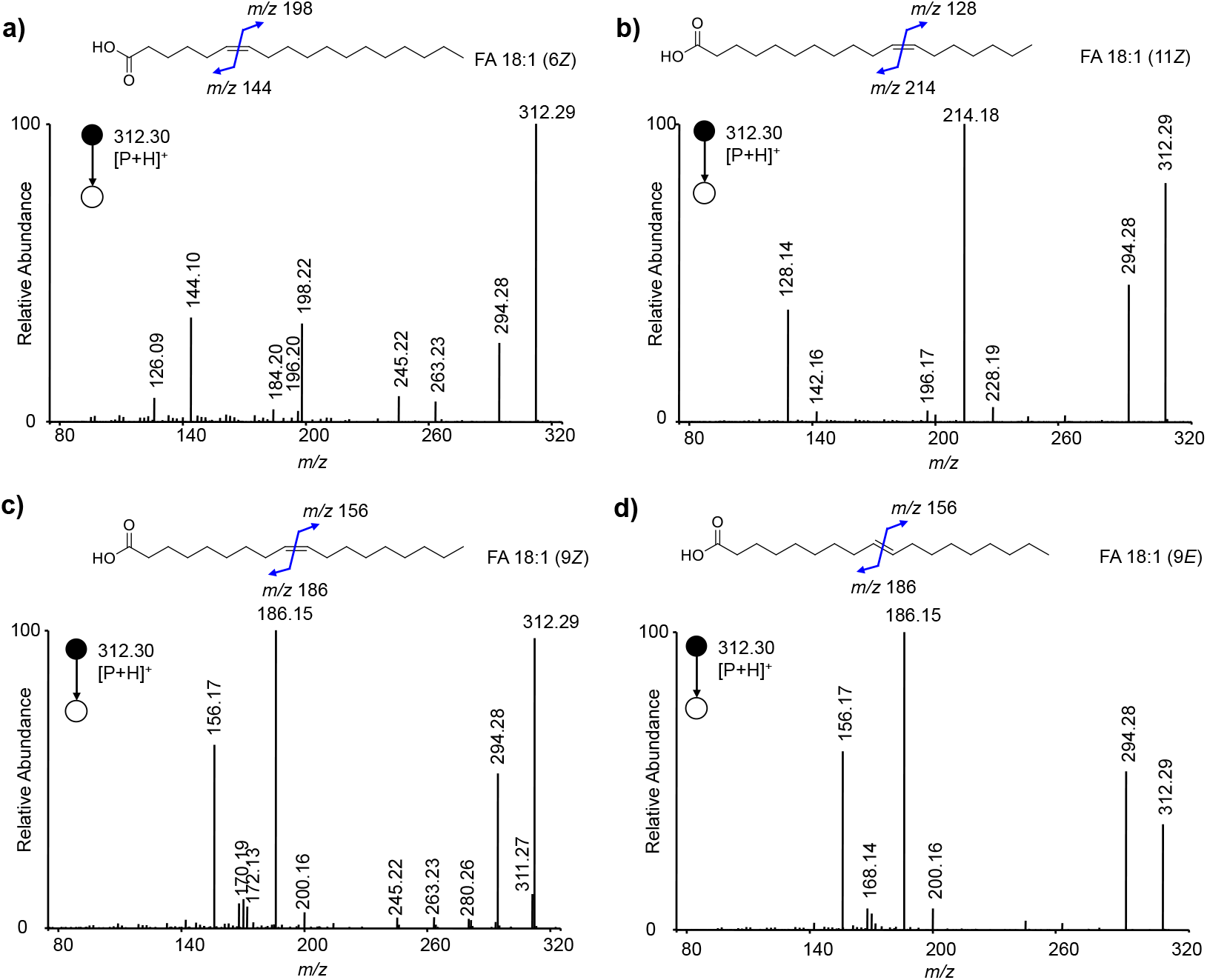
MS/MS spectra of the aziridination product ions formed by the reaction of TsONHCH_3_ with fatty acids. **a)**-**d)** MS/MS spectra of the product ions derived from FA 18:1 (6*Z*), FA 18:1 (11*Z*), FA 18:1 (9*Z*) and FA 18:1 (9*E*). The chemical structures of the lipids show the positions of C=C bonds and the *m/z* values of the possible diagnostic fragment ions of the aziridination reaction products *via* CID.

### Investigation of the aziridination reaction with polyunsaturated fatty acids

Polyunsaturated fatty acids are lipids with multiple C=C bonds, and the precise identification of the position of each C=C bond is a challenge. To verify the feasibility of using this aziridination reaction to identify the double bond positions in polyunsaturated fatty acids, we used a variety of polyunsaturated fatty acids to try them out. The experimental results showed that the aminating reagent TsONHCH_3_ was able to react with them all efficiently, resulting in the generation of an aziridine structure at the double bond position. Interestingly, even though the aliphatic chains contain multiple unsaturated double bonds, addition to one of the double bonds accounts for the majority of the reaction products. For example, for FA 18:2 (9*Z*, 12*Z*), we can see four characteristic fragment ions containing nitrogen positive ions (*m/z* 154.16, 186.15, 114.13 and 226.18) in MS/MS spectrum, derived from two single addition reaction products of aziridination (Figure 4a). Likewise, the positions of C=C bonds in FA 18:2 (9*Z*, 11*E*), FA 18:2 (9*E*, 12*E*), and FA 18:3 (9*Z*, 12*Z*, 15*Z*) were identified correctly by the characteristic fragments of the their aziridination products (Figure 4b-4d). More importantly, even the fatty acid containing four C=C bonds [FA 20:4 (5*Z*, 8*Z*, 11*Z,* 14*Z*)] could be identified accurately. The fragments for assigning each of the C=C bond could be detected in the MS/MS spectrum (Figure 4e).

**Figure 4.**
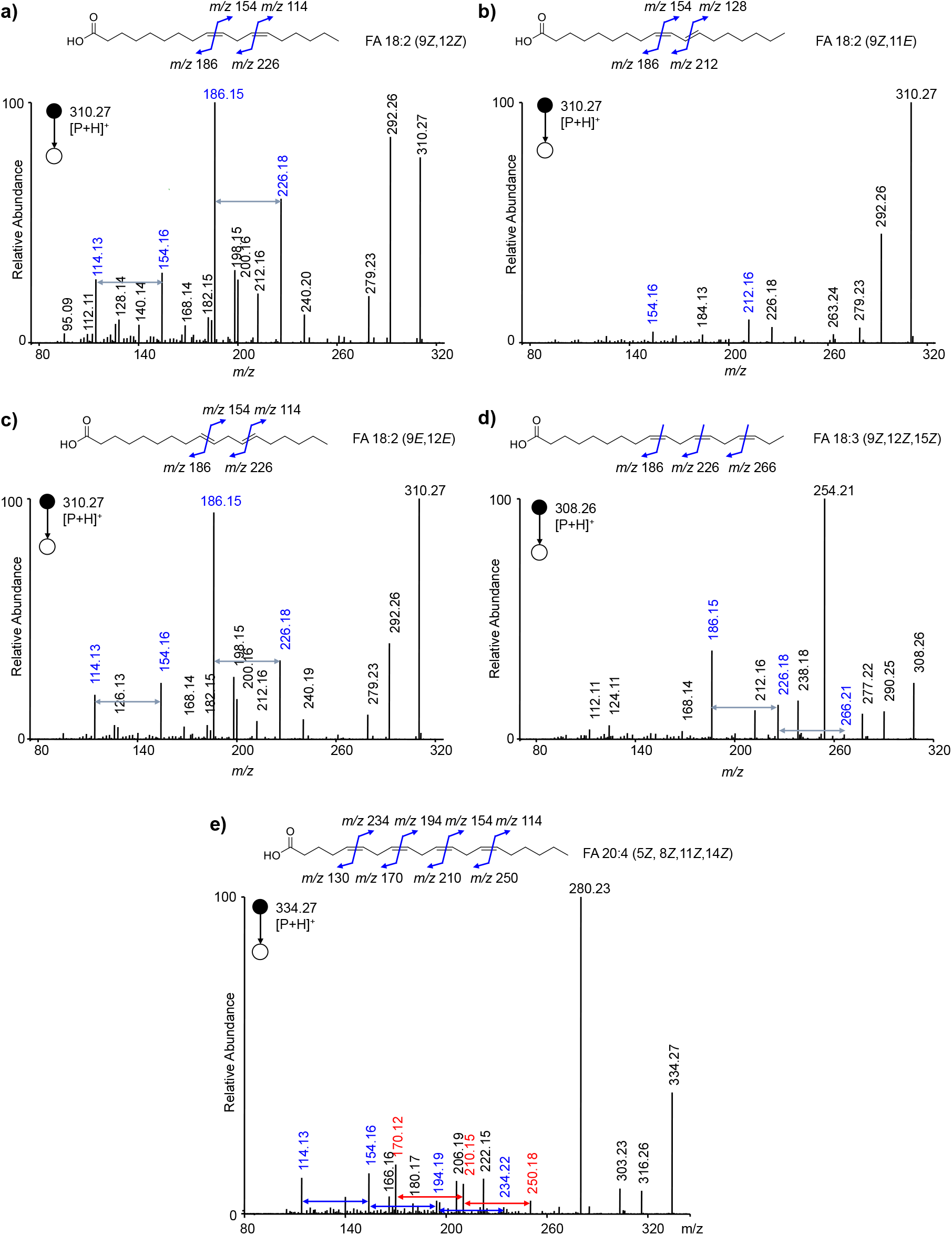
MS/MS spectra of the aziridination product ions formed by the reaction of TsONHCH_3_ with polyunsaturated fatty acids. **a)**-**e)** MS/MS spectra of the product ions derived from FA 18:2 (9*Z*, 12*Z*), FA 18:2 (9*Z*, 11*E*), FA 18:2 (9*E*, 12*E*), FA 18:3 (9*Z*, 12*Z*, 15*Z*), and FA 20:4 (5*Z*, 8*Z*, 11*Z*, 14*Z*). The chemical structures of the lipids show the positions of C=C bonds and the *m/z* values of the possible diagnostic fragment ions of the aziridination reaction products *via* CID.

### Identification of the locations of C=C bonds in glycerophospholipids

To explore the versatility of this aziridination reaction on the identification of the locations of C=C bonds, different kinds of glycerophospholipids (GPLs) such as phosphatidylglycerol (PG), phosphatidylethanolamine (PE), phosphatidylserine (PS), phosphatidyl acid (PA) and phosphatidylcholine (PC) were tested (Figure 5 and 6). Different from the analysis of fatty acids, the head groups of GPs were easily lost in CID, and produce the predominant fragment ions with the loss of these head groups. The losses of phospholglycerol, phosphorylethanolamine, phosphoserine, phosphoric acid and phosphocholine in aziridination products of GPs were the dominant competitive fragmentation pathways compared with that of the aziridine structure. Thus, the characteristic fragment ions that indicating the position of C=C bond would be obtained in muti-stage MS/MS (MS^3^) from the precursor ions losing the head groups (Figure 5 and 6). For instance, in the MS^3^ spectrum of protonated aziridination product of PG 18:0/18:1 (9*Z*), the characteristic ions at *m/z* 479.41 and 508.42 derived from the MS^2^ fragment ion at *m/z* 634.58 (by losing the phosphoglycerol head group) were detected that denote the position of C=C bond at Δ9 (Figure 5a). The fragment ions at *m/z* 312.29 and 294.28 correspond to the ions of aminating fatty acyl chain (FA 18:1) and its derived ion by losing a water, respectively. These two ions could provide additional information for the structural analysis of the lipid isomers. These results indicate the good compatibility of this aziridination reaction to GPLs and the corresponding resolving power for C=C bond positions.

**Figure 5.**
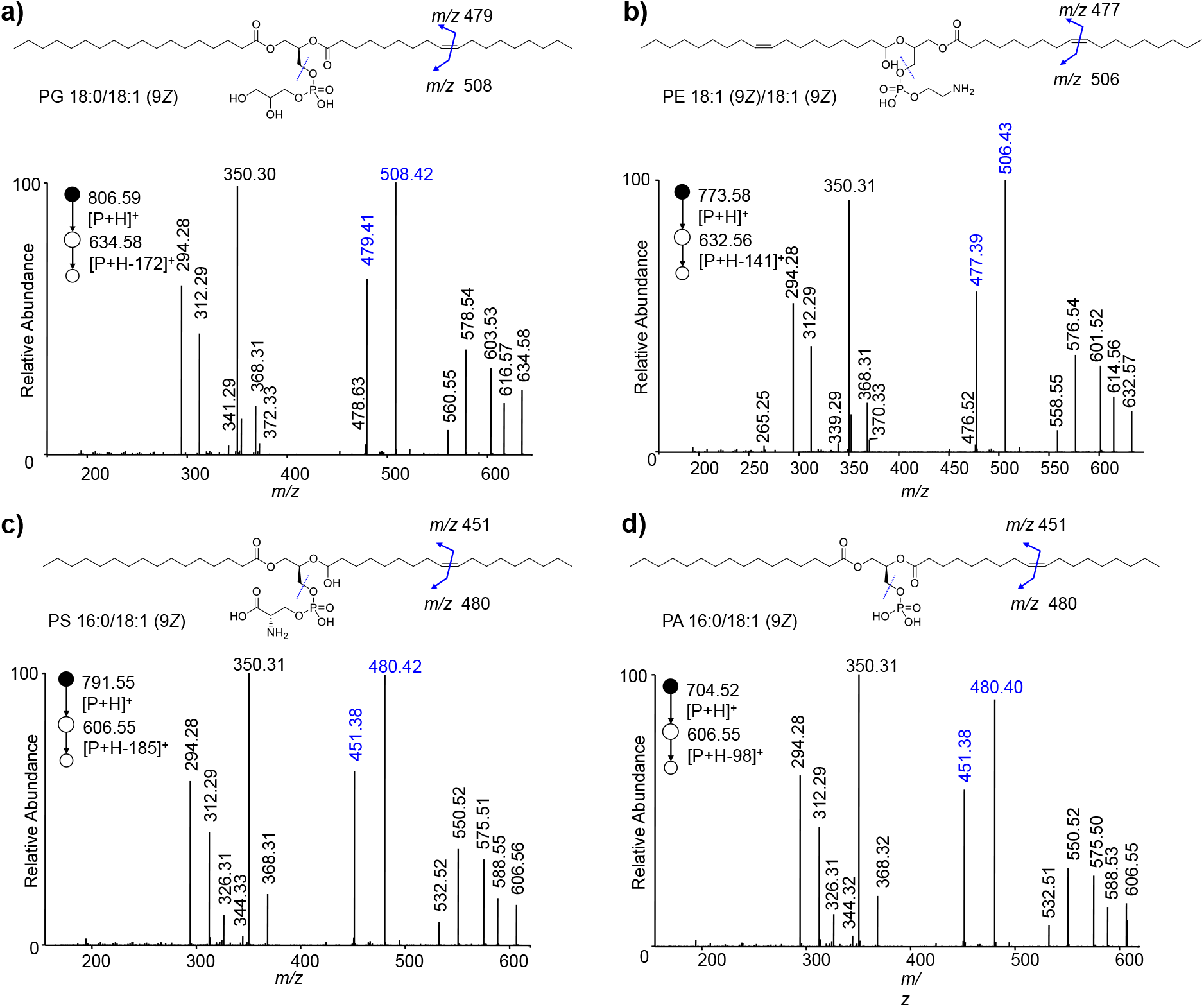
MS^3^ spectra of the aziridination product ions formed by the reaction of TsONHCH_3_ with glycerophospholipids. **a)**-**d)** MS^3^ spectra of the product ions derived from PG 18:0/18:1 (9*Z*), PE 18:1 (9*Z*)/18:1 (9*Z*), PS 16:0/18:1 (9*Z*), and PA 16:0/18:1 (9*Z*). The chemical structures of the lipids show the positions of C=C bonds and the *m/z* values of the possible diagnostic fragment ions of the aziridination reaction products *via* CID.

**Figure 6.**
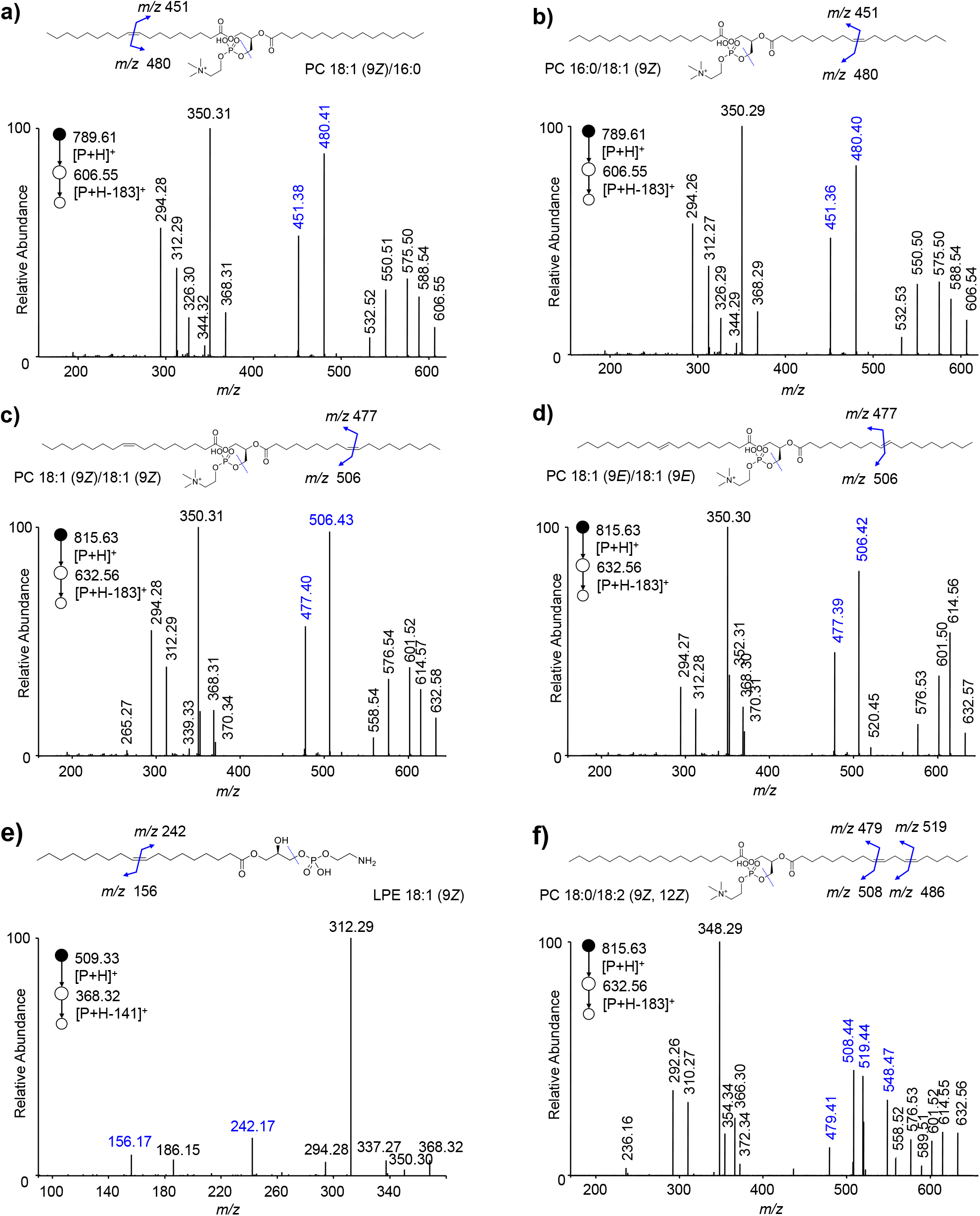
MS^3^ spectra of the aziridination product ions formed by the reaction of TsONHCH_3_ with phosphatidylcholines and lysophosphatidylethanolamine. **a)**-**e)** MS^3^ spectra of the product ions derived from PC 18:1 (9*Z*)/16:0, PC 16:0/18:1 (9*Z*), PC 18:1 (9*Z*)/18:1 (9*Z*), PC 18:1 (9*E*)/18:1 (9*E*), LPE 18:1 (9*E*), and PC 18:0/18:2 (9*Z*, 12*Z*). The chemical structures of the lipids show the positions of C=C bonds and the *m/z* values of the possible diagnostic fragment ions of the aziridination reaction products *via* CID.

### Investigating the versatility of this aziridination reaction with different unsaturated lipids

This aziridination reaction were also applied to revolve the C=C bond positions in glycerides. As shown in Figure 7a, the fragment of the aminating product of diacylglycerol (DG) 16:0_18:1 (9*Z*) containing the nitrogen positive ion was observed at *m/z* 498.42 in the MS/MS spectrum. The fragmentation pathways were showed in Figure 7b. For triacylglycerol (TG) 18:1 (9*Z*)/18:1 (9*Z*)/18:1 (9*Z*), the fragment ion at *m/z* 788.67 was detected in the MS/MS spectrum after the CID of the aziridination product, which indicates the C=C bond at Δ9 for each fatty acyl chain (Figure 7c). For the polyunsaturated TG, both two characteristic fragment ions at *m/z* 784.64 and 824.68 could be detected clearly that show the unambiguous locations of two double bonds at Δ9 and Δ12 in TG 18:2 (9*Z*, 12*Z*)/18:2 (9*Z*, 12*Z*)/18:2 (9*Z*, 12*Z*) (Figure 7d). More importantly, the N-Me aziridination products for each double bond could be high efficiently separated by LC, and gave clear MS/MS spectra for each product to identify the position of C=C bonds (Figure S1 in Supporting Information). These results indicate the compatibility of this approach in the identification of C=C bond positional isomers for polyunsaturated glycerides.

**Figure 7.**
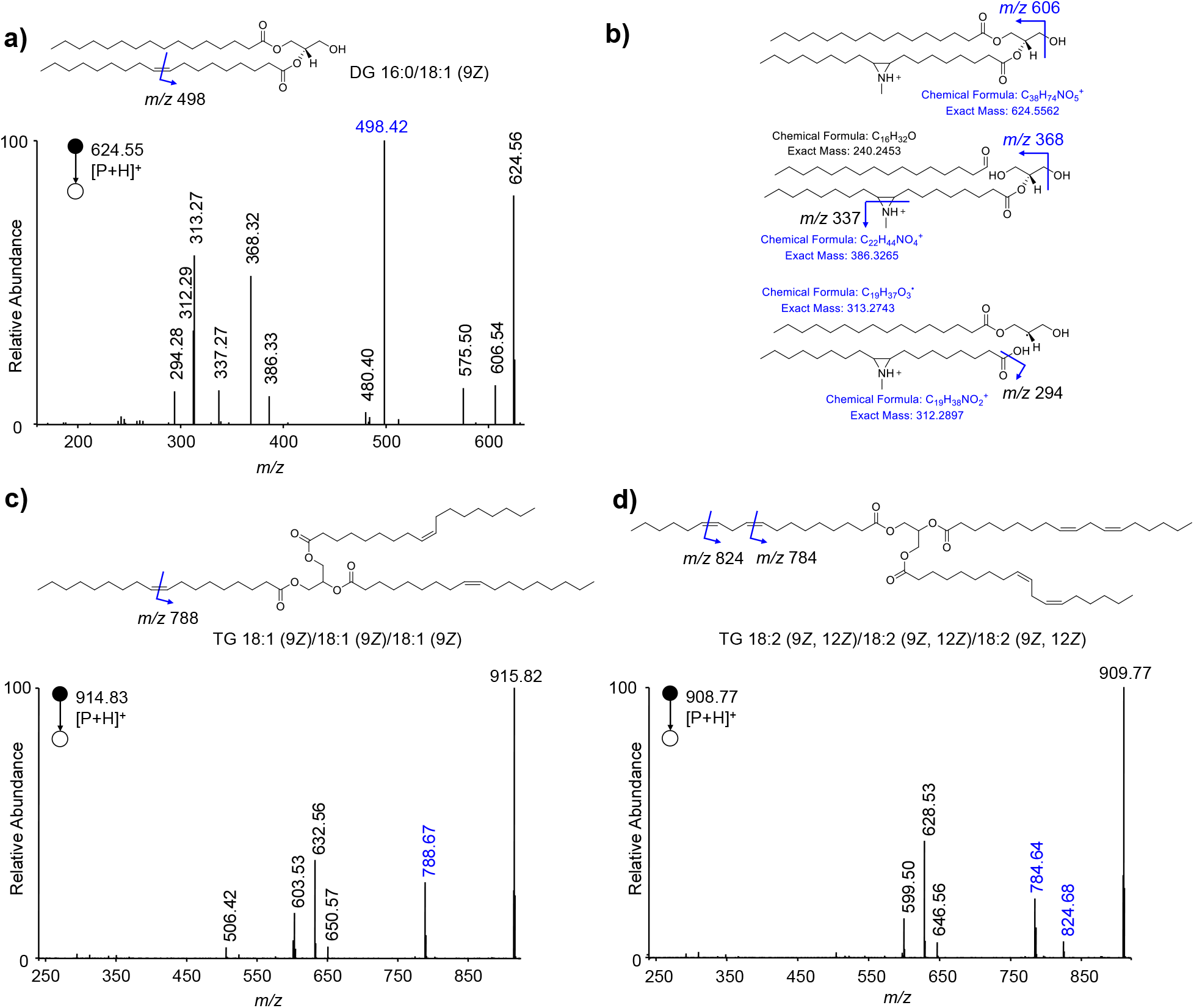
MS/MS spectra of the aziridination product ions formed by the reaction of TsONHCH_3_ with glycerides. **a)-b)** MS/MS spectra of the product ions derived from **a)** DG 16:0/18:1 (9*Z*) and **b)** its possible fragmentation pathways. **c)-d)** MS/MS spectra of the product ions derived from TG 18:1 (9*Z*)/18:1 (9*Z*)/18:1 (9*Z*) and TG 18:2 (9*Z*, 12*Z*)/(9*Z*, 12*Z*)/(9*Z*, 12*Z*). The chemical structures of the lipids show the positions of C=C bonds and the *m/z* values of the possible diagnostic fragment ions of the aziridination reaction products *via* CID.

In addition, the identification of C=C bond in sphingolipids was also conducted (Figure S2 in Supporting Information). Sphingolipids are a class of lipids containing a backbone of sphingoid bases, a set of aliphatic amino alcohols that includes sphingosine. The representative sphingolipids are ceramides (Cer) and sphingomyelins (SM). Ceramides are *N*-acylated sphingoid bases lacking additional head groups, whereas sphingomyelins have a phosphocholine molecule with an ester linkage to the 1-hydroxy group of a ceramide. Like the analysis of GPLs, the phosphocholine head group of SM is easily to be lost in CID (Figure S2a). Therefore, the characteristic fragment ions denoting the position of C=C bond in the fatty acyl chain of SM d18:1/16:1 (11*E*) could be observed in MS^3^ spectrum (Figure S2b). For SM d18:1/12:0 that doesn’t have the unsaturated fatty acyl chain, the fragment ion indicating the double bond position in sphingoid base was detected at *m/z* 226.25 in MS^3^ spectrum (Figure S2c). However, the same fragment ion could be observed clearly in MS^2^ spectrum for Cer d18:1/16:0, because there is no easily fragmented phosphocholine head group in it (Figure S2d).

### Quantitative analysis of lipid C=C location isomers

As mentioned before, the aziridination reaction products of unsaturated lipids have a tertiary amine structure with a very high proton affinity, which could greatly improve the sensitivity in the identification of unsaturated lipids based on this chemical derivatization strategy. To further examine the quantitative capability of this method, the absolute quantification of the method for unsaturated lipids was first investigated. Taking methyl oleate as an example (Figure 8a), the intensity of both two characteristic fragment ions (**F1** and **F4** at *m/z* 156 and 200) of its aminating products were proportional to the concentration of methyl oleate, showing good linearities in the range of 25-500 nM (*R*^2^ = 0.998), and the detection limit could reach to 1 nM.

**Figure 8.**
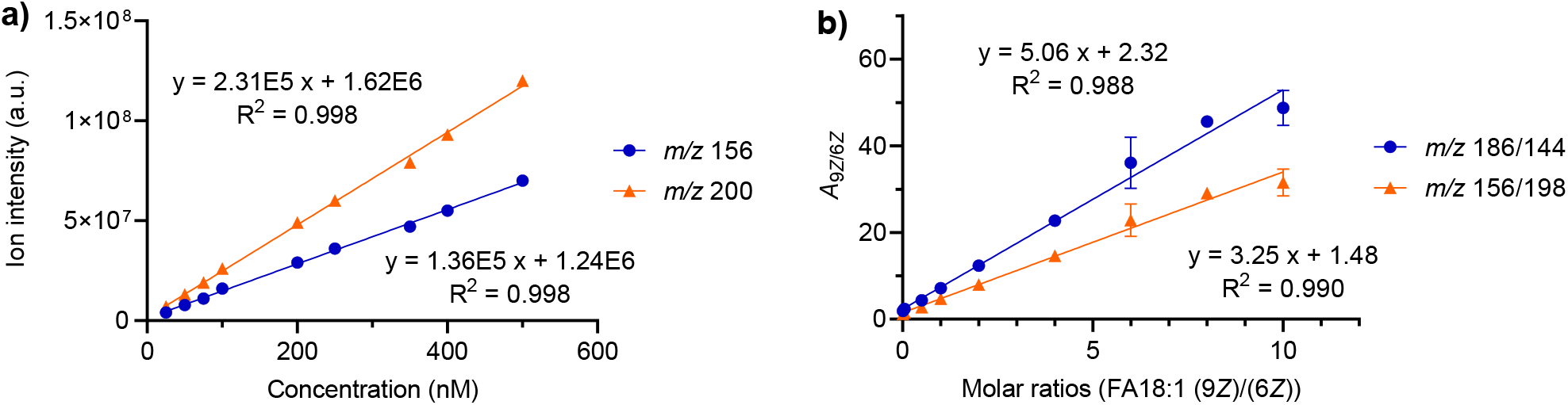
Quantitative analysis of lipid C=C location isomers. **a)** Calibration curves for quantitation of methyl oleate in the range of 25-500 nM. **b)** Linear relationship established between the peak area ratio (A_9*z*_/A_6*z*_) of the diagnostic ions and molar ratio (C_9*z*_/C_6*z*_) of the two FA 18:1 C=C location isomers.

Quantification of the ratio of the C=C isomers was performed with a series of mixtures of FA 18:1 (9*Z*) and FA 18:1 (6*Z*) isomer. As shown in Figure 8b, the ratios of the peak areas of the same type of diagnostic ions from each isomer (i.e., **F1**-type fragments at *m/z* 186 and 144, and **F4**-type fragments at *m/z* 156 and 198, derived from 9*Z* and 6*Z*, respectively), were found to be directly proportional to the molar ratios of these two isomers with the total concentration kept constant (0.5 μM). The good linearities (R^2^ = 0.988 and 0.990) were obtained with molar ratios from 1:100 to 10:1. These results demonstrated the good quantitative capability of the proposed aziridination reaction-based method for lipid C=C isomers.

### Application of the aziridination derivatization in structural lipidomics

To validate the aziridination reaction in structural lipidomics, the human serum sample was used for the structurally resolving of the lipid structures at C=C positional isomer level. As shown in Figure 9, three steps were taken for identifying the lipids with the information of C=C bond positions. First, the lipids in serum sample were extracted and analysed by LC-MS/MS. The mass data was analyzed with MS-DIAL software including extracting peaks and matching the MS and MS/MS data with that in LipidBlast library to generate a list of lipids with the RT, formula, extract mass, mass error, adduct type and MS/MS ions. Then, on the basis of the identified lipids in human serum, another extended unsaturated lipid list was obtained including the information of derived lipids (by the mass increase of 30 Da during the aziridination, [**P**+H]^+^), and certain putative unsaturated lipids that were missed in the lipidomics analysis. Finally, the lipid extract after the derivatization reaction was subjected to targeted tandem mass spectrometric analysis based on the parent ions in extended lipids list. By analyzing the characteristic fragment ions of the derived lipids, the locations of the C=C bonds in unsaturated lipids could be obtained.

**Figure 9.**
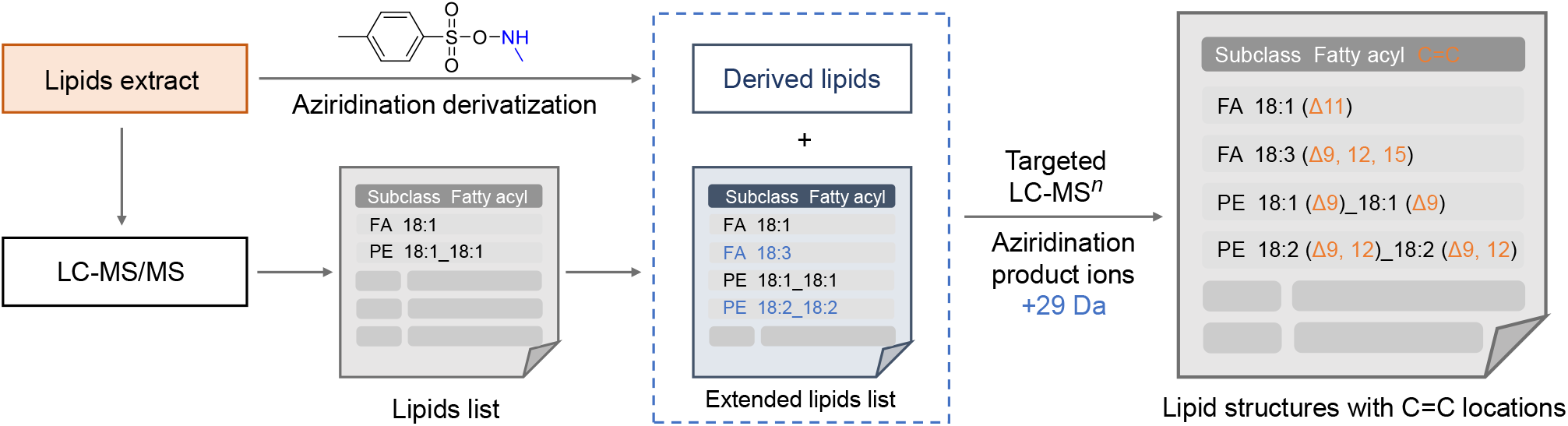
Workflow for the analysis of the lipids in serum extract based on the aziridination reaction and LC-MS.

The glycerol phospholipid was analyzed with MS^3^, and the precursor ions for MS^3^ was calculated by subtracting the mass of the head group (PE 141, PC 183, PG 172, PS 189, PA 98, PI 26) from the mass of the protonated derivative product. Other lipids were analyzed with MS^2^. We found that the diagnostic ion **F1** (or the ion that losing the head group from **F1**) has higher intensity than the ion **F4** (or the ion that losing the head group from **F4**). Thus, the **F1**-type ions were typically used for the identification of the location of C=C bonds, and they could also be calculated by subtracting the mass of **F2** ion from the mass of the product ion (**P**). Then, the C=C bond positions can be readily deduced one by one with *de novo* analysis. With the developed workflow, 505 lipids in serum were identified including 36 saturated lipids, in which 425 of them were further identified at C=C location level. Among, 113 pair of mono-unsaturated and 4 pair of poly-unsaturated C=C bond positional isomers were identified (Supplementary Datasheet).

We also observed that the muti-stage MS/MS (MS^3^) acquisition method for the analysis of GPLs have several advantages. First, the MS^3^ spectra are very clean with less noise ions, and the diagnostic ions for locating the C=C bond have high intensity comparing with other product ions. Benefit from this advantage, 126 of PCs were identified with the C=C bond locations (including 34 of ether-linked PCs). In addition, the fragment ions in the low mass range of the MS^3^ spectra could indicate the structures of the aminating fatty acyl chains, which can be used to easily distinguish isomer mixture. Take the isomer of PC 32:1 (PC 14:0_18:1 and PC 16:0_16:1) as an example, the product ion at *m/z* 284 could be detected in the MS^3^ spectrum for PC 16:0_16:1, but the product ion at *m/z* 312 was detected in the MS^3^ spectrum for PC 14:0_18:1. Obviously, the fragment ion at *m/z* 284 denoting the fatty acyl chain FA 16:1 after the aziridination reaction, and fragment ion at *m/z* 312 indicates the existence of FA 18:1 fatty acyl chain. For the analysis of DG and TG, the fragment ions of the aminating fatty acyl chains in the low mass range of the MS/MS spectra could also facilitate the resolving of the isomers. The cholesterol esters (CE) are special lipids that hard to be ionized in electrospray ionization. In this study, the highly efficient aziridination of C=C bonds was observed in the fatty acyl chains of CE. It is worth mentioning that 37 of CE were detected in serum (Supplementary Datasheet).

## Experimental section

### Lipid nomenclature

The shorthand notations for lipids structural annotations were adopted from LIPID MAPS. For example, TG 16:0/18:0/18:1 (9*Z*) indicates triacylglycerol (TG) lipid containing one C16 and two C18 fatty acyl chains. The number “0” and “1” after the colon represent the degree of unsaturation of each fatty acyl chain. For the position of the carbon-carbon double bond, it is indicated by the number in the braket (*n*). The number *n* indicates the ordinal of carbon starts from the alpha carbon at the carboxyl end of the aliphatic chain, and the carbon-carbon double bond is located between the nth and (*n*+1)th carbons. The capital letters “*Z*/*E*” following the position of the double bond indicate its *“trans/cis”* configurations. Forward slash (“/”) suggests that the *sn-* position is known and the *sn*-1 chain is placed before “/”, while a underscore (“_”) means that *sn*-position is unspecified.

### Materials

Fatty acids including FA 18:1 (9*Z*), FA 18:1 (9*E*), FA 18:1(6*Z*), FA 18:1 (11*Z*), FA 18:2 (9*Z*, 11*E*), FA 18:2 (9*Z*, 12*Z*), FA 18:2 (9*Z*, 12*Z*,15*Z*) and FA 20:4 (5*Z*, 8*Z*, 11*Z*, 14*Z*) were purchased from Sigma-Aldrich. Sphingolipids including Cer d18:0/12:0, SM d18:0/16:1 and SM d18:0 /16:1, glycerides including DG 16:0/18:1 (9*Z*), TG 18:1 (9*Z*)/18:1 (9*Z*)/18:1 (9*Z*), TG 18:2 (9*Z*, 12*Z*)/18:2 (9*Z*, 12*Z*)/18:2 (9*Z*, 12*Z*), TG 16:0/18:1 (9*Z*)/18:2 (9*Z*,12*Z*), and glycerol phospholipids including PG 18:0/18:1 (9*Z*), PS 16:0/18:1 (9*Z*), PA 18:0/18:1 (9*Z*), PE 18:1 (9*Z*)/18:1 (9*Z*), PC 18:1 (9*Z*)/18:1 (9*Z*), PC 18:1 (9*E*)/18:1 (9*E*), PC 18:0/18:2 (9*Z*, 12*Z*), PC 16:0/18:1 (9*Z*) and PC 18:1 (9*Z*)/16:0 were purchased from Avanti Polar Lipids (Alcbaster, AL, USA). Other solvents were purchased from Innochem (Beijing, China) and meet or exceed the analytical grade standard.

### Lipid extraction and sample preparation

The human serum was collected at Renmin Hospital of Wuhan University. The protocol of fatty acid extraction from human serum is described below: 400 μL methyl tert-butyl ether (MTBE) was added into the 1.5 mL Eppendorf tube which contains serum samples 100 μL, and subjected to vortex for 30 s and ultrasonic mixing for 10 min in ice water. After vortex and sonicate, Then, 80 μL MeOH and 100 μL H_2_O were added successively with the same steps as above. After the vortex and sonicate, the mixture was centrifuged for 10 min at 10,000 rpm to separate phases. The upper MTBE phase was taken to a new EP tube and dried with nitrogen flow. The dried lipid extracts were redissolved with 100 μLTFE solvent and subjected to injection with mass spectrometry analysis or to aziridination reaction.

### Derivatization and analysis

For the aziridination derivatization of lipid standards or serum extract, the solutions were dried with nitrogen flow and then redissolved with 1 mL of trifluoroethanol (TFE) solution that contains the aminating reagent TsONHCH_3_ (final concentration 10 mM) and catalyst Rh_2_(esp)_2_ (final concentration 1 mM). The reaction mixture was vortexed in thermostatic oscillator for 60 min. After the reaction, the solution was centrifuged at 16,000 g for 10 min, and the supernatant was collected to do the LC-MS analysis.

### Liquid chromatographic and mass spectrometric analysis

All data except for the regular lipidomics analysis of human serum were accomplished with LTQ-Orbitrap Elite mass spectrometer (Thermo Scientific, Germany). The main mass parameters were set up as below: sheath gas (N_2_): 60 arbitrary units; auxiliary gas (N_2_): 20 units; capillary temperature: 380°C; spray voltage: 3.8 kV; collision energy: 40 V; the resolution of full mass scan, 30,000 and The MS/MS scan was induced by collision dissociation (CID), with a resolution of 15,000.

A Waters ACQUITY UPLC BEH C18 Column (2.1 mm × 100 mm, 1.7 mm) was performed by the UltiMate 3000 UPLC system (DIONEX, Thermo Scientific, Germany) and maintained at 40°C for separation of lipid species. The flow rate was set as 0.3 *μ*L/min. Gradient elution consisted of mobile phase A (H_2_O), mobile phase B[(ACN/H_2_O, 60:40, v/v) and C [(IPA/ACN, 90:10, v/v). All mobile phase were mixed with 10 mM NH_4_OAc and 0.1% formic acid. The optimal chromatographic gradient program were achieved with three mobile phase as below: 50%A-50%B, 0 min; 100%B,10 min, holding for 3 min; 70%B-30%C, 14 min; 48%B-52% C, 16 min; 37%B-63%C, 18 min; 32%B-68%C, 19 min; 27%B-73%C, 34 min.

### Data analysis

MS-DIAL version 4.60 was used for the process of LC-MS data for human serum lipid extract analysis. The raw data must be transformed into Abf. files. More details were presented at http://prime.psc.riken.jp/compms/msdial/main.html. Thermo Xcalibur mass spectrometry data system was used for extracting target ion, analyzing the targeted secondary ion spectrum, and calculating its peak area.

## Supporting information

Supplementary Datasheet

## Acknowledgements

This work was financially supported by the National Key Research and Development Program of China [2021YFC2700700 (S.M.C.)] and the National Natural Science Foundation of China [22074111 (S.M.C.), 22004092 (G.F.F) and 22004093 (Q.Q.W.)]. We also thank the support of the start-up funds of Wuhan University (S.M.C.) and the National Youth Talents Plan of China (S.M.C.).

