## Supporting Information for "Direct N-Me Aziridination Reaction Enables Pinpointing C=C Bonds in Lipids with Mass Spectrometry"

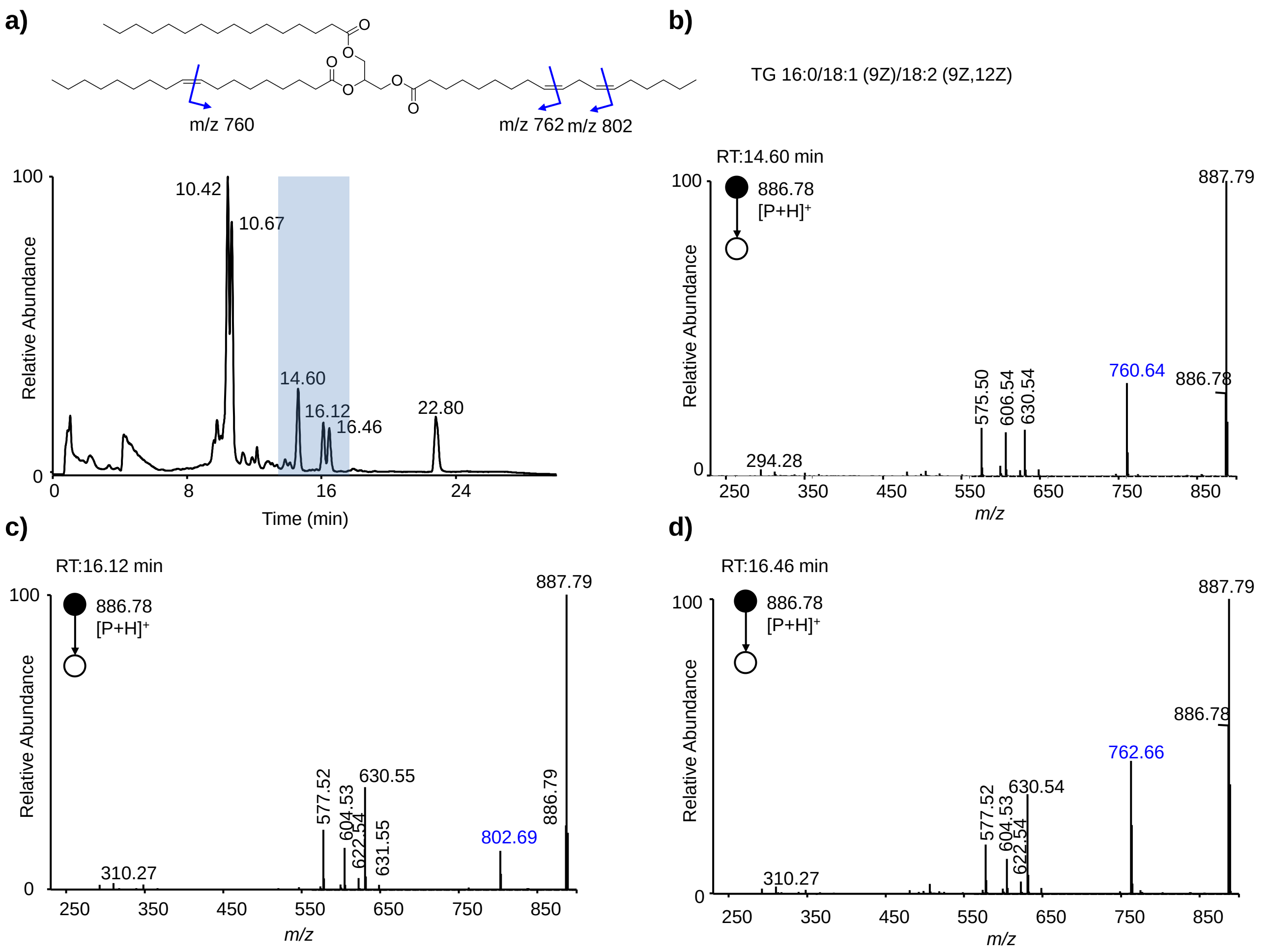


**Figure S1.** Analysis of the location of C=C bonds in TG 16:0/18:1 (9*Z*)/18:2 (9*Z*, 12*Z*). **a)** Base peak chromatogram of reaction solution of TsONHCH$_3$ with TG 16:0/18:1 (9*Z*)/18:2 (9*Z*, 12*Z*). **b)-d)** MS/MS spectra of the aziridination product ions derived from the products eluted at **b)** 14.60, **c)** 16.12, and **d)** 16.46 min in LC. The chemical structures of the lipids show the positions of C=C bonds and the *m/z* values of the possible diagnostic fragment ions of the aziridination reaction products *via* CID.

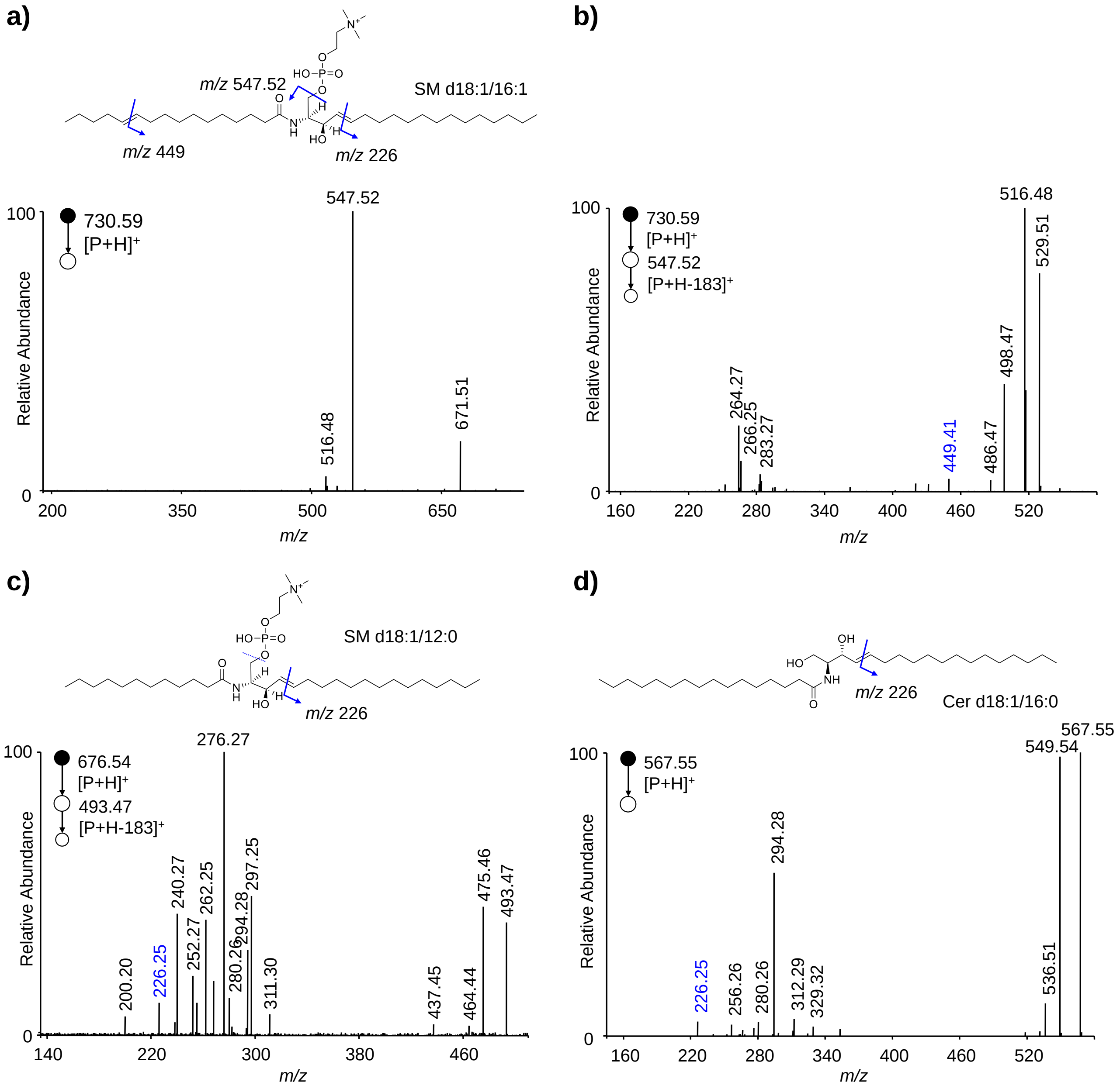


**Figure S2.** MS$^2$ and MS$^3$ spectra of the aziridination product ions formed by the reaction of $TsONHCH_3$ with sphingolipids. **a)-b) a)** MS$^2$ and **b)** MS$^3$ spectra of the product ions derived from SM d18:1/16:1 (11*E*). **c)** MS$^3$ and spectrum of the product ions derived from SM d18:1/12:0. d) MS$^2$ spectrum of the product ions derived from Cer d18:1/16:0. The chemical structures of the lipids show the positions of C=C bonds and the *m/z* values of the possible diagnostic fragment ions of the aziridination reaction products *via* CID.

**Synthesis of the $TsONHCH_3$:**

**Tert-butyl (tosyloxy)carbamate:** tert-butyl hydroxycarbamate (13.3 g, 100 mmol) and TsCl (19.3 g, 105 mmol) was added into a 1000 mL round bottom flask, then added $Et_2O$ (400 mL). $Et_3N$ (14.1 mL, 105 mmol) was added dropwise to the flask at 0 °C. The mixture was stirred at 0 °C for 20 minutes, then warmed to room temperature and stirred for 4 hours. The reaction mixture was filtered through celite and the filter cake was rinsed with $Et_2O$. The filtrate was concentrated in vacuo to give the crude product, and the resulting solid was taken up in n-hexane (200 mL) and stirred at room temperature for 30 minutes. The mixture was filtered and the filter cake was rinsed with hexane. The resulting white solid was dried in vacuo to obtain the title compound. **$^1$H NMR** (600 MHz, Chloroform-*d*) δ 7.89 (d, *J* = 8.3 Hz, 2H), 7.58 (s, 1H), 7.37 (d, *J* = 8.4 Hz, 2H), 2.47 (s, 3H), 1.30 (s, 9H). **$^{13}$C NMR** (151 MHz, Chloroform-*d*) δ 154.1, 146.0, 130.5, 129.7, 129.7, 83.9, 27.7, 21.8.

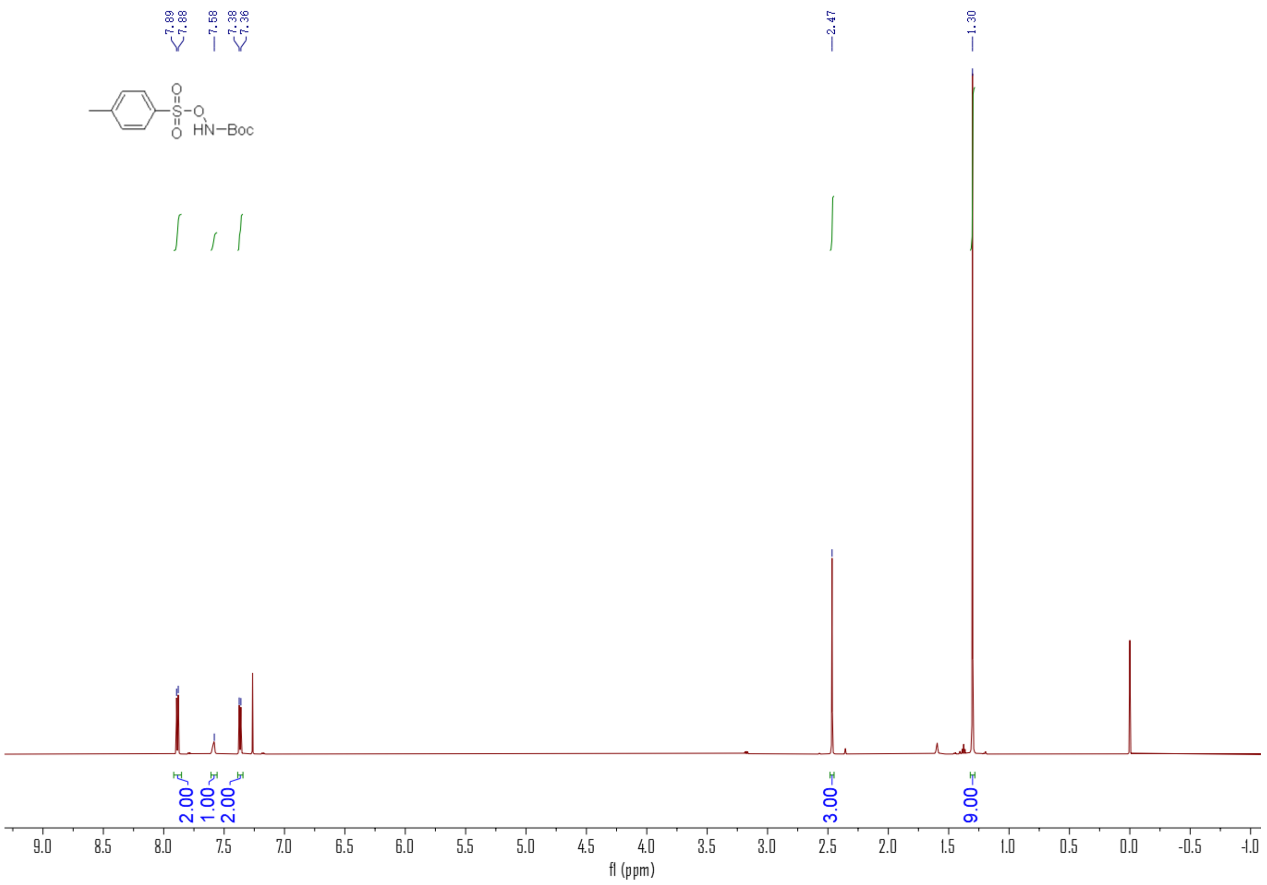

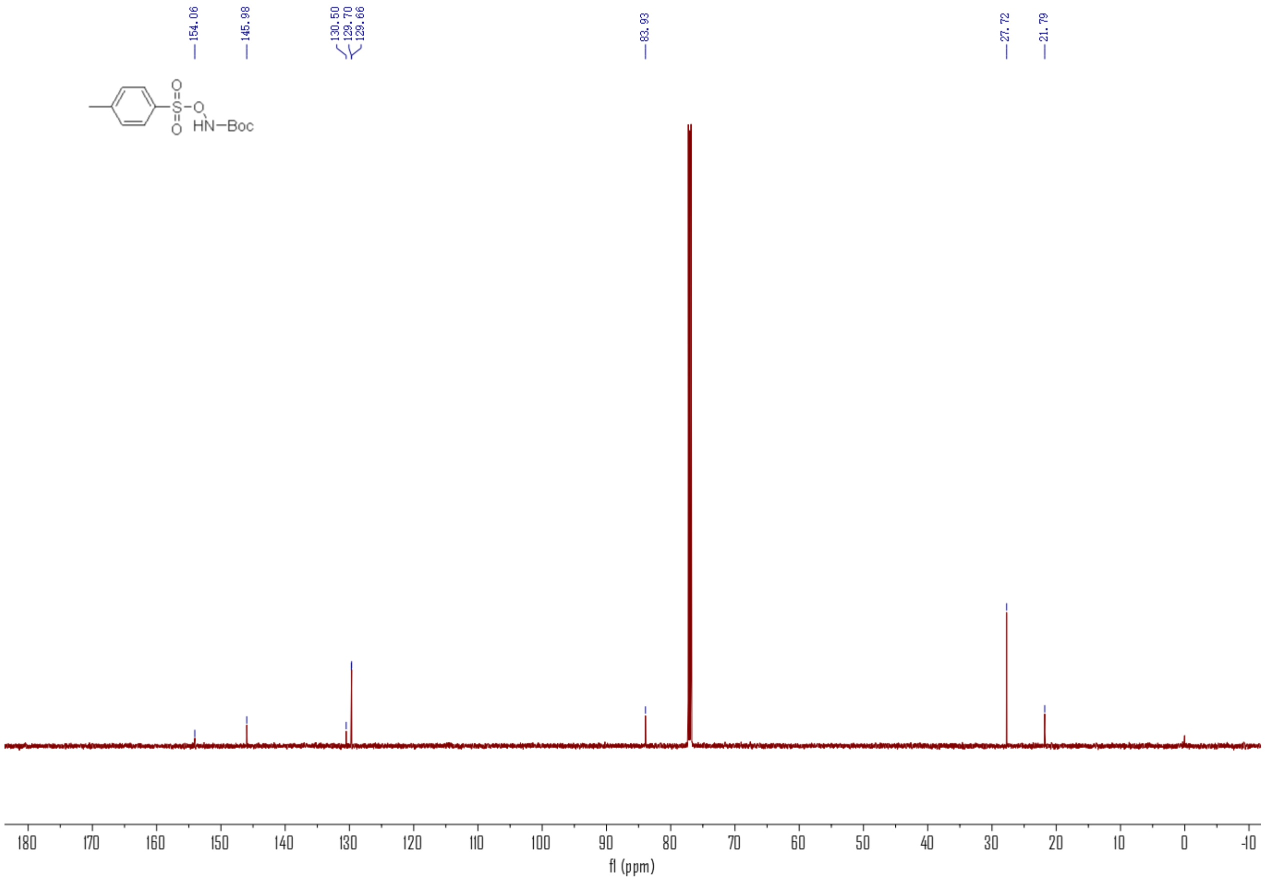


**Tert-butyl methyl(tosyloxy)carbamate:** TsONHBoc (2.87 g, 10 mmol), $PPh_3$ (2.75 g, 10.5 mmol) was added into a 100 mL round bottom flask, then added 50 mL dry THF. Methanol (0.34g, 10.5 mmol) was added under nitrogen. Added DIAD (2.12 g, 10.5 mmol) dropwise in an ice bath. The mixture was stirred at room temperature overnight. Purification by silica gel chromatography using PE/EA as elution solvent gave tert-butyl methyl(tosyloxy)carbamate as a white solid (2.71 g, 90%).

**Tert-butyl (tosyloxy)carbamate:** tert-butyl methyl(tosyloxy)carbamate (2 g, 6.67 mmol) in 6 mL DCM solution was added to TFA (9.86 mL, 13.34 mmol ) in an ice bath. The mixture was stirred at 0 ºC for 3 h. TFA and DCM were then removed under reduced pressure in vacuo. The concentrated oil was dissolved in 50 mL DCM, and the solution was washed with saturated sodium bicarbonate solution, dried over anhydrous sodium sulfate and concentrated under reduced pressure, obtained white solid (1.12 g, 84%). **$^1$H NMR** (600 MHz, Chloroform-*d*) δ 7.85 (d, $J$ = 8.3 Hz, 2H), 7.34 (d, $J$ = 7.7 Hz, 2H), 5.44 (s, 1H), 2.74 (s, 3H), 2.45 (s, 3H). **$^{13}$C NMR** (151 MHz, Chloroform-*d*) δ 145.0, 132.2, 129.6, 129.0, 40.2, 21.7.

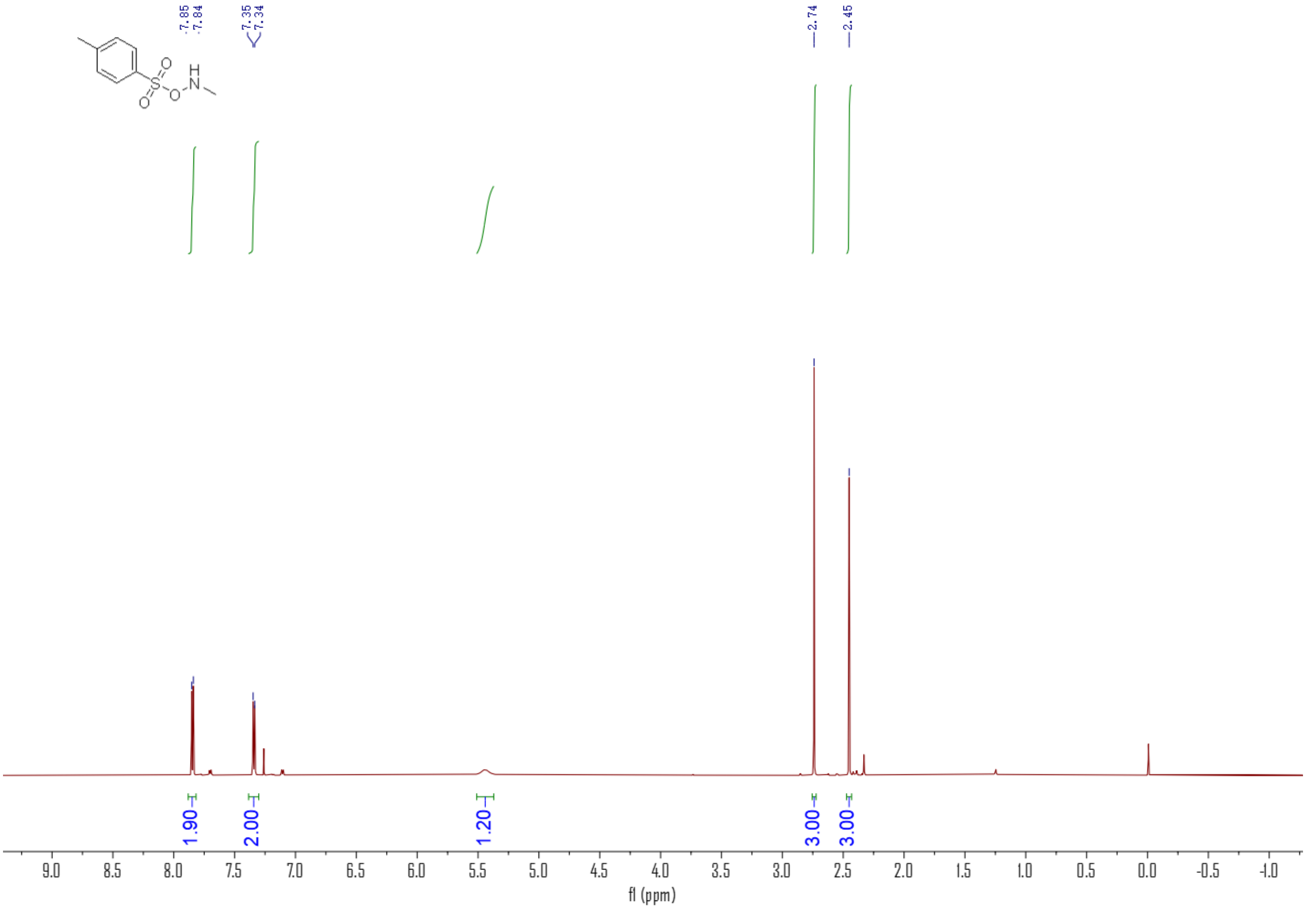

7.85
7.84
7.35
7.34
2.74
2.45
1.90
2.00
1.20
3.00
3.00
f1 (ppm)


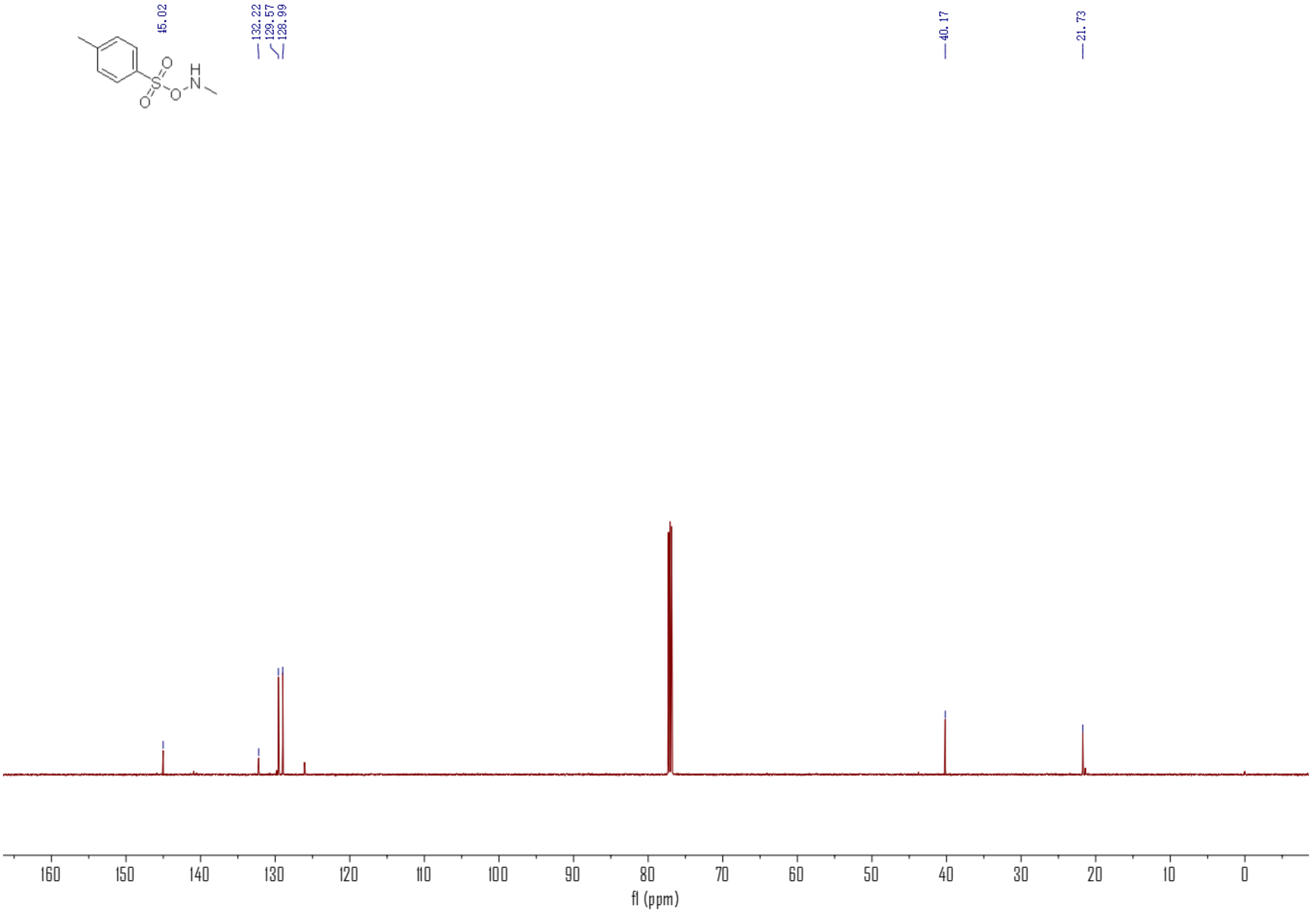

45.02
132.22
129.57
128.99
40.17
21.73
f1 (ppm)
